# Tolloid cleavage activates latent GDF8 by priming the pro-complex for dissociation

**DOI:** 10.1101/154823

**Authors:** Viet Q. Le, Roxana E. Iacob, Yuan Tian, William McConaughy, Justin Jackson, Yang Su, Bo Zhao, John R. Engen, Michelle Pirruccello-Straub, Timothy A. Springer

## Abstract

Growth differentiation factor 8 (GDF8)/Myostatin is a latent TGF­β family member that potently inhibits skeletal muscle growth. Here, we compared the conformation and dynamics of precursor, latent, and Tolloid­cleaved GDF8 pro­complexes to understand structural mechanisms underlying latency and activation of GDF8. Negative stain electron microscopy (EM) of precursor and latent pro­complexes reveals a V­shaped conformation that is unaltered by furin cleavage and sharply contrasts with the ring­like, cross­armed conformation of latent TGF­β1. Surprisingly, Tolloid­cleaved GDF8 does not immediately dissociate, but in EM exhibits structural heterogeneity consistent with partial dissociation. Hydrogen–deuterium exchange was not affected by furin cleavage. In contrast, Tolloid cleavage, in the absence of prodomain–growth factor dissociation, increased exchange in regions that correspond in pro-TGF-β1 to the α1-helix, latency lasso, and β1 strand in the prodomain and to the β6’–7’ strands in the growth factor. Thus, these regions are important in maintaining GDF8 latency. Our results show that Tolloid cleavage activates latent GDF8 by destabilizing specific prodomain–growth factor interfaces and primes the growth factor for release from the prodomain.

## Introduction

Growth and differentiation factor 8 (GDF8, myostatin) is one of 33 members of the transforming growth factor-β (TGF-β) family, which in addition to GDFs and TGF-βs includes activins, inhibins, and bone morphogenetic proteins (BMPs). GDF8 potently negatively regulates skeletal muscle development. Mutation of GDF8 in humans and multiple animal species results in a hypermuscular and low body fat phenotype (Clop, 2006, Grobet, Martin et al., 1997, McPherron, Lawler et al., 1997, McPherron & Lee, 1997, Schuelke, Wagner et al., 2004). Clinical trials with GDF8 antagonists aim to treat muscle wasting associated with muscular dystrophy, cancer cachexia, sarcopenia, trauma, diabetes, and chronic obstructive pulmonary disease (Bogdanovich, 2002, Cohen, Nathan et al., 2014, Smith, 2013).

Like other members of the TGF-β family, GDF8 is synthesized as a proprotein precursor consisting of a signal peptide, a large N-terminal prodomain, and a smaller C-terminal growth factor (GF) domain (Fig. 1A). In the endoplasmic reticulum (ER), the signal peptide is removed, the GDF8 monomers dimerize, and disulfide bonds form, including one between the two GF moieties, thus yielding the inactive pro-complex precursor (pro-GDF8). After secretion into muscle tissue, a proprotein convertase (PC) cleaves between the prodomain and growth factor domain, to yield latent GDF8 (Anderson, Goldberg et al., 2008).

**Figure 1.**
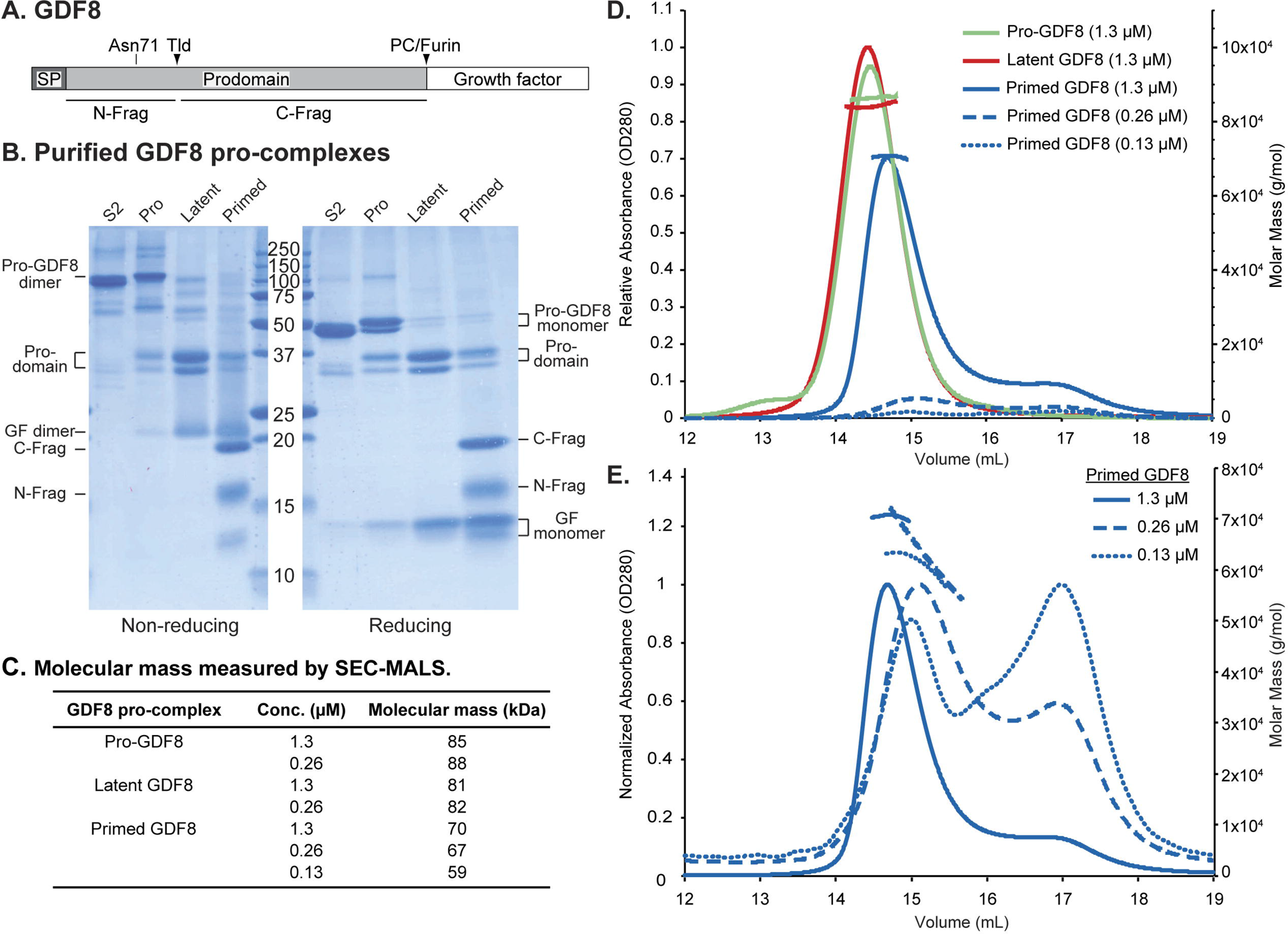
Expression and characterization of GDF8 pro-complexes. **A.**Schematic diagram of the GDF8 proprotein shown to scale. The N-glycan attachment site at Asn48 and the Tolloid (TLD) and Proprotein convertase (PC)/furin cleavage sites are indicated above the diagram. The boundaries of the N-terminal prodomain fragment (N-Frag) and C-terminal prodomain fragment (C-Frag) generated by TLD cleavage are indicated below the diagram. SP = signal peptide. **B.** Coomassie blue-stained SDS-PAGE gels under non-reducing (left) and reducing (right) conditions of purified S2-produced pro-GDF8 (S2) and mammalian expressed pro-GDF8 (Pro), latent GDF8 (Latent), and primed GDF8 (Primed). GF = growth factor. Expected molecular mass of the disulfide-linked pro-GDF8 = 81.8 kD, pro-GDF8 monomer = 40.9 kD, Prodomain = 28.5 kD, TLD-cleaved N-terminal prodomain fragment (N-Frag) = 9.5 kD, TLD-cleaved C-terminal prodomain fragment C-Frag = 19 kD, growth factor (GF) dimer = 24.8 kD, and growth (GF) monomer = 12.4 kD. **C–E.** Characterization of GDF8 pro-complexes by size exclusion chromatography coupled to multi-angle light scattering (SEC-MALS). **C.** Table summarizing the calculated molecular mass of pro-, latent, and primed GDF8 as determined by SEC-MALS. **D.** Comparison of SEC elution profiles and molecular masses of pro-, latent, and primed GDF8. The left Y-axis indicates relative absorbance (OD280) and the right Y-axis indicates calculated molar mass (g/mol). **E.** Comparison of SEC elution profiles of primed GDF8 at 1.3, 0.26, and 0.13 μM. The peak absorbance for each sample was normalized to 1. The left Y-axis indicates normalized absorbance (OD280) and the right Y-axis indicates calculated molar mass (g/mol).

Pro-complexes of GDF8, its close relative GDF11, and TGF-βs 1, 2, and 3 are latent as recombinant proteins (Ge, Hopkins et al., Gentry, Webb et al., 1987, Hill, Davies et al., 2002, Khalil, 1999, Lee & McPherron, 2001, Thies, Chen et al., 2001, Wakefield, Smith et al., 1988, Zimmers, Davies et al., 2002). After cleavage by PCs between the prodomain and GF, the prodomain and GF remain noncovalently associated in all characterized family members to date; however, the strength of this association differs (Hinck, Mueller et al., 2016). TGF-β family GFs are typically active at pM to nM concentrations. In contrast, the GF-prodomain dissociation constants are often higher. For example, the BMP9 GF has an EC50 for cellular activation of ~ 1 nM but the prodomain only inhibits with an IC50 of ~ 100 nM and has a Kd for the GF in a similar range; therefore, at bioactive concentrations the GF and prodomain are largely dissociated and pro-complexes show similar activity to the isolated GF in bioassays (Mi, Brown et al., 2015). Whereas isolated pro-complexes of most TGF-β family members are active, the GDF8 prodomains and GF dimer remain tightly non-covalently associated in a latent pro-complex that is not competent for signaling (Hill et al., 2002, Lee & McPherron, 2001, Thies et al., 2001); the prodomains block receptor binding to the GF and prevent activation of downstream signaling (Lee & McPherron, 2001).

Why some TGF-β family members are active and others are latent as pro-complexes is incompletely understood. The basis for latency of pro-TGF-β1 was illuminated by its crystal structure, which showed a ring-like structure with the GF dimer surrounded by the prodomain dimer (Shi, Zhu et al., 2011). On one side, large arm domains in each prodomain disulfide link to one another and interact with the GF near the periphery of the ring, with a solvent-filled channel between the prodomain and GF at the center of the ring. On the other side, smaller prodomain helical and latency lasso structural elements form a straitjacket that wraps around the GF. Cysteines near the N-terminus of the TGF-β1 prodomain disulfide link in the ER to “milieu molecules” that store latent pro-TGF-β1 on cell surfaces or in the extracellular matrix for subsequent activation (Springer, Dong et al., 2016). Other family members including GDF8 have similar cysteines that might link to milieu molecules and further stabilize latency; however, their structural and functional relevance is unclear. More recent structures of two non-latent TGF-β1 family members, BMP9 and activin-A, reveal important differences among pro-complexes (Mi et al., 2015, Wang, Fischer et al., 2016). Both have V-shaped or linear overall shapes, termed open-armed, that contrast with the ring-like, cross-armed conformation of TGF-β1. Further differences were revealed, particularly with BMP9, in the manner of association of straitjacket elements with the GF.

Here, we ask why GDF8 is latent, and what changes when it becomes activated. GDF8 and its close relative GDF11 are activated by BMP1/Tolloid (TLD) metalloprotease-mediated cleavage of the prodomain between the straitjacket elements and the arm domain (Ge et al., 2005, Wolfman, McPherron et al., 2003). Tolloid-like protein 2 (TLL2), used in this paper, is among the most active on GDF8 of the four TLD proteases found in mammals (Wolfman et al., 2003), and is the only TLD protease expressed in muscle (Scott, Imamura et al., 2000). While TLD cleavage clearly activates signaling by the GF, whether the two prodomain fragments rapidly dissociate from the GF after cleavage, or remain associated with the GF in a “primed” state, is not known. Here, we compare pro-GDF8, the state prior to PC cleavage; latent GDF8, the state after PC cleavage; and primed GDF8, a state after TLD cleavage in which we found the persistence of substantial prodomain-GF association. We use two orthogonal techniques, negative stain electron microscopy (EM) and hydrogen-deuterium exchange mass spectrometry (HDX-MS) to compare these three states. The results provide important insights into the structure and mechanism of activation of GDF8. Our results and those of two other groups were concurrently submitted to BioRxiv (Cotton, Fischer et al., 2017, Le, Iacob et al., 2017, Walker, McCoy et al., 2017b). After submission, we were able to read one another’s manuscripts and learn of remarkable concordance. In this MS, we refer not only to what we initially deduced based on comparisons to structures of previously published TGF-β1 family member pro-complexes (Le et al., 2017), but also to what we learned from concurrently submitted crystal structure and small-angle X-ray scattering analysis of pro-GDF8 (Cotton et al., 2017, Walker et al., 2017b).

## Results

### Molecular composition of three types of GDF8 prodomain – GF complexes and their functional activity

To interrogate the effect of pro-complex maturation by PC cleavage and activation by Tolloid cleavage, we generated three different GDF8 pro-complexes: 1) the uncleaved GDF8 precursor pro-complex (pro-GDF8), 2) furin-cleaved GDF8 pro-complex (latent GDF8), and 3) furin- and Tolloid-cleaved GDF8 pro-complex (primed GDF8). Stable 293 transfectants were grown in the presence (to generate pro-GDF8) or absence (to generate partially PC-cleaved GDF8) of the PC inhibitor decanoyl-Arg-Val-Lys-Arg-chloromethylketone. Pro-GDF8 or partially PC-cleaved material was purified from conditioned media by Ni-NTA and size-exclusion chromatography (SEC). To obtain latent GDF8 and primed GDF8, partially PC-cleaved GDF8 was incubated with purified furin protease alone or with furin and TLL2-conditioned media, respectively. FLAG-tagged proteases were then removed on an affinity column and pro-complexes were purified by an additional round of SEC.

Purified pro-GDF8 was predominantly uncleaved and migrated as a 110 kDa disulfide-linked precursor dimer under non-reducing conditions and a doublet of 52 and 47 kDa precursor monomers under reducing conditions (Fig. 1B). Pro-GDF8 produced in S2 insect cells, shown for comparison, migrated at 100 and 47 kDa in non-reducing and reducing SDS-PAGE, respectively. S2 cells make high mannose N-glycans. The identical migration of the insect cell material to the 47 kDa band from mammalian 293 cells in reducing gels suggests that the 52 and 47 kDa mammalian pro-GDF8 monomers have complex and high-mannose glycans, respectively (Anderson et al., 2008); pro-GDF8 has a single predicted N-linked glycan attachment site at Asn-71 in the straitjacket region (Fig. 1A).

Furin cleavage of pro-GDF8 to create latent GDF8 converted the precursor dimer of 110 kDa in non-reducing SDS-PAGE to 35 and 31 kDa prodomain monomers and a 20 kDa GF dimer (Fig. 1B). Reducing SDS-PAGE of latent GDF8 showed the same 35 and 31 kDa prodomain monomer bands as non-reducing PAGE together with a 13 kDa GF monomer.

TLL2 cleavage to obtain primed GDF8 converted the prodomain monomer bands of 35 and 31 kDa seen in latent GDF8 to bands at 19 kDa and 15 kDa under both non-reducing and reducing conditions (Fig. 1B). Bands from reducing SDS-PAGE were subjected to Edman degradation. The N-terminal sequence of the 15 kDa band was HXXXXXNEN, corresponding to the N-terminal sequence of the His-tagged prodomain (HHHHHHNEN). The N-terminal sequence of the 19 kDa band was DXSXXGXLE, showing that it corresponds to the fragment C-terminal to the TLD-cleavage site (DDSSDGSLE). Additional bands corresponding to a 20 kDa GF dimer and 13 kDa GF monomer were seen in non-reducing and reducing SDS-PAGE, respectively. Since the primed GDF8 had run as a symmetric peak in SEC after TLL2 cleavage, these results showed that the N-terminal and C-terminal prodomain fragments largely remained associated with the GF dimer, and justified terming this material primed GDF8. We further term the N-terminal 15 kDa and C-terminal 19 KDa cleaved prodomain fragments the N-prodomain and C-prodomain fragments, respectively. The greater diffuseness in SDS-PAGE of the N-prodomain than C-prodomain fragment and higher observed mass in SDS-PAGE of 15 kDa compared to the expected protein mass of 9.5 kDa of the N-prodomain fragment correlate with its predicted N-linked glycan attachment site. The mass from SDS-PAGE of the C-prodomain fragment of 19 kDa agreed with its predicted protein mass of 19 kDa. The mass from SDS-PAGE of 13 kDa for the GF monomer agreed with its predicted protein mass of 12.4 kDa. Moreover, a mass of 20 kDa for the GF dimer in non-reducing SDS-PAGE is consistent with the expectation that intra-chain disulfide bonds in each monomer increase migration of the GF dimer in SDS-PAGE.

The molecular masses of the three classes of GDF8 pro-complexes made in mammalian cells were characterized by SEC with multi-angle light scattering (SEC-MALS), which estimates the total mass of glycoproteins independently of their shape (Fig. 1C-E). Samples were applied at concentrations ranging from 0.13 to 1.3 μM. Pro-GDF8 and latent GDF8 had molecular masses of 86 and 82 kDa, respectively, consistent with their expected protein masses of 82 kDa and additional N-glycosylation (Fig. 1C-D). Furthermore, similar masses were estimated for pro-GDF8 and latent GDF8 when they were used at differing concentrations of 1.3 and 0.26 µM (Fig. 1C).

In contrast, primed GDF8 yielded a main peak with a substantially lower mass of 70 to 59 kDa and a secondary, later-eluting peak. As the concentration of primed GDF8 in the experiment was reduced from 1.3 to 0.26 to 0.13 µM, the main peak in gel filtration eluted later with a decrease in estimated molecular mass (Fig. 1C and E), and the amount of material in the secondary peak increased relative to the main peak (Fig. 1E). These results show concentration-dependent complex dissociation; and thus that Tolloid cleavage primes GDF8 for dissociation. The samples had been stored frozen and diluted just prior to SEC-MALS evaluation; therefore, dissociation of prodomain fragments from the GF dimer occurred on the time scale of sample handling and SEC-MALS. Moreover, the concentration dependence of the molecular mass of the main peak and the increasing proportion of the secondary peak with decreasing primed GDF8 concentration showed that rebinding of dissociated prodomain fragments to the GF occurred and suggested dissociation constants in the range of experimentally used concentrations. Notably, the highest concentration of primed GDF8 used in the experiments had a calculated molecular weight that was less than that of pro- or latent GDF8. To achieve a mass of 70 kDa, it appears that the GF dimer must be present in the complex and associate with a combined total of ~3 prodomain fragments on average. Overall, the results suggest that cleavage at the TLD site enables partial dissociation of the primed GDF8 pro-complex in a concentration-dependent fashion.

We compared the signaling activities of the three GDF8 pro-complexes to one another and to recombinant, commercially obtained GDF8 using GDF8-responsive luciferase reporter cells (Fig. 2). Because of the high signaling potency of GDF8, GDF8 complexes were diluted to much lower concentrations than in the SEC-MALS experiments. Pro-GDF8 had little or no activity. Latent GDF8 induced minimal signaling with an estimated EC50 of 2.62 nM, consistent with previous observations and likely reflecting the dissociation of the prodomain at such low protein concentrations (Hill et al., 2002, Lee & McPherron, 2001, Thies et al., 2001). In contrast, primed GDF8 and the purified GDF8 growth factor signaled equivalently, with EC50 values of 0.074 and 0.078 nM, respectively. The equivalent activities of primed GDF8 and the GDF8 GF showed that TLL2 cleavage completely activated latent GDF8 and resulted in a 35-fold higher signaling potency relative to latent GDF8.

**Figure 2.**
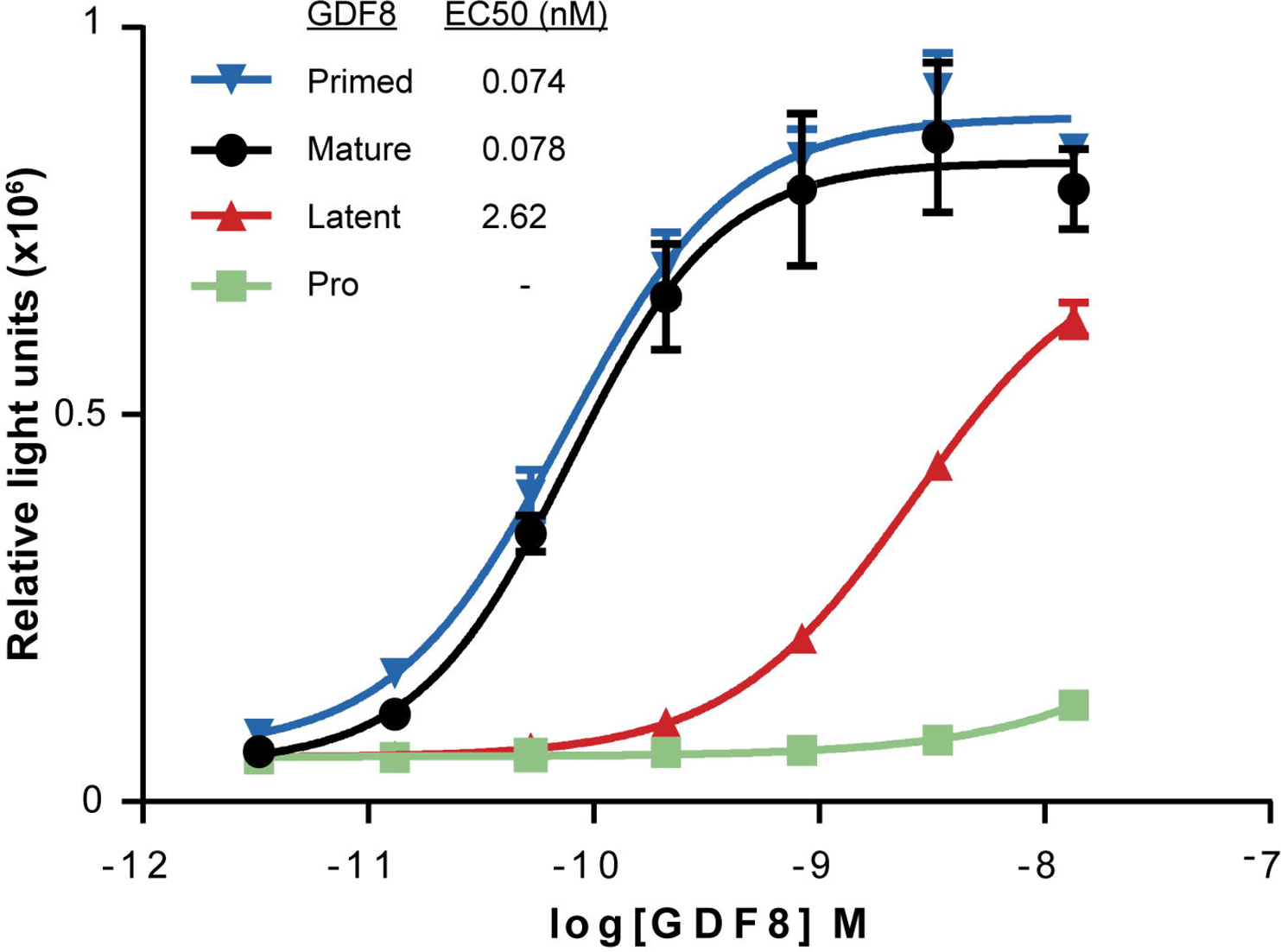
Signaling activity of GDF8 pro-complexes compared to mature GDF8 growth factor as measured in a cell-based, Smad-responsive luciferase reporter assay.

### HDX-MS

We used HDX-MS to investigate the structural differences among the three types of GDF8 pro-complexes, and more specifically, to gain insights into the effects of PC and TLL2 cleavage on polypeptide backbone dynamics (Fig. 3). For HDX-MS, GDF8 pro-complexes at concentrations of 11–25.4 μM were diluted 15-fold into 99% D_2_O buffered at pD 7.5 and incubated for time periods varying from 10 s to 4 h to allow exchange of pro-complex backbone amide hydrogens with solvent deuteriums. Labeling was quenched in H_2_O buffered at pH 2.5 with tris (2-carboxyethyl) phosphine (TCEP) to reduce disulfides. Samples were digested online with pepsin and subjected to liquid chromatography coupled to mass spectrometry (LC-MS) to determine, for each peptide, the number of hydrogens substituted with deuterium. Measurements were in triplicate, i.e., each time point was analyzed three times in an independent measurement. Overall, we obtained ~88% peptide coverage for each of the three GDF8 pro-complexes (Fig. 3A and S1 and S2). Coverage of the prodomain was essentially complete; less coverage of the GF correlated with its high content of disulfide-bonded cysteines which challenges reduction.

**Figure 3.**
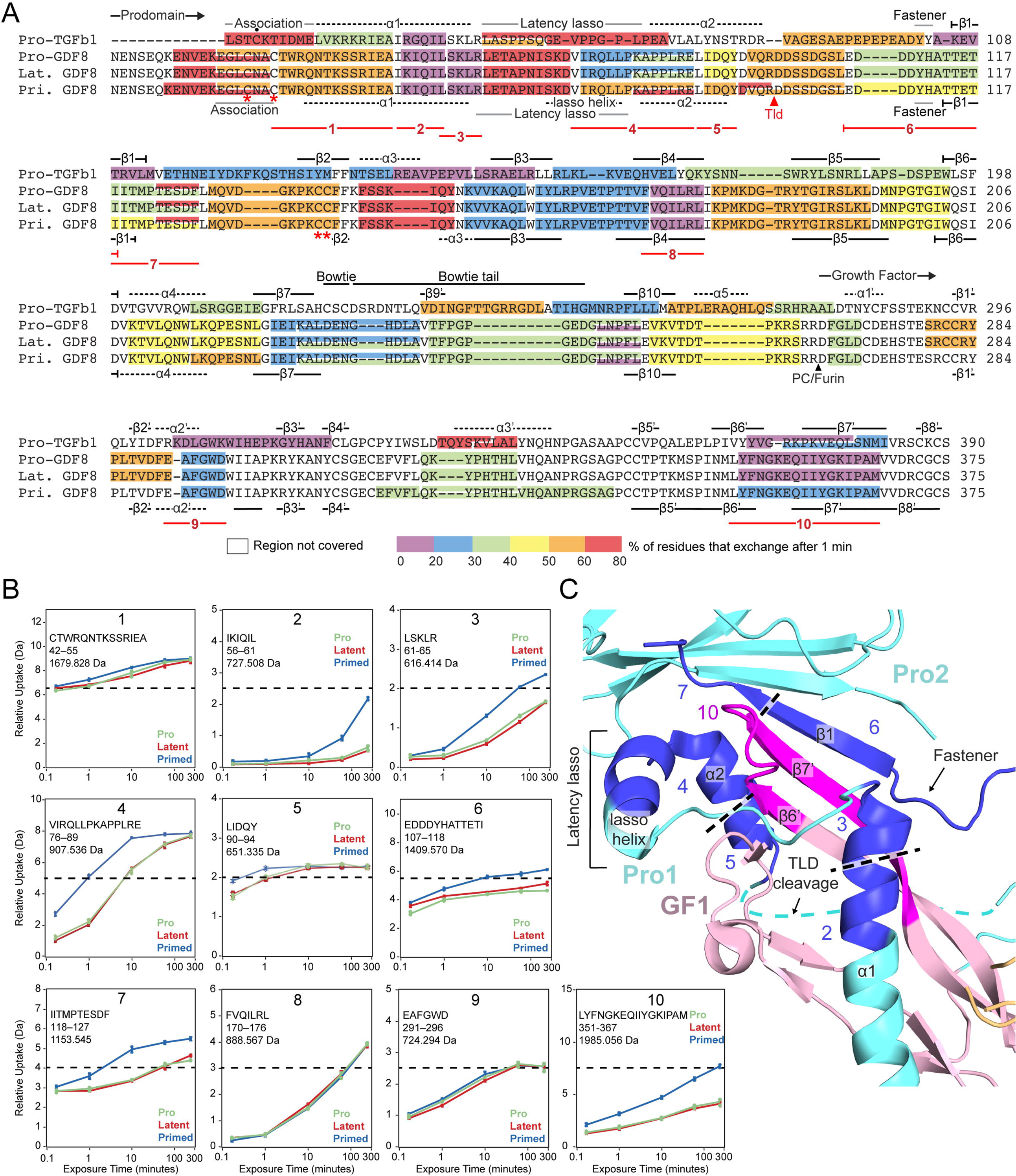
HDX-MS of GDF8 pro-complexes at pD 7.5. **A.**HDX of GDF8 pro-complexes compared to pro-TGF-β1 R249A. The color (according to the scale shown) represents the percentage of deuterium incorporation for each peptide after 1 min in deuterium and are superimposed onto a structure-based sequence alignment of TGF-β1 R249A and GDF8. Secondary structures based on the pro-TGF-β1 and pro-GDF8 crystal structure are shown above and below the sequences, respectively. PC/furin and TLD-cleavage sites are indicated by arrowheads. Dot (•) marks the Cys4 residue in pro-TGF-β1 that disulfide links to LTBPs and GARP. Asterisks (*) mark free cysteines in the GDF8 prodomain. Numbered lines below the alignment represent identical peptides 1–9 followed by HDX-MS from pro-, latent, and primed GDF8 samples that are compared in the text and in panels B & C. **B.** Deuterium incorporation graphs for peptic peptides 1–9 from pro-, latent, and primed GDF8. Values represent the mean of three independent measurements; error bars, s.d. **C.** Peptides that reveal enhanced HDX in primed GDF8 were mapped onto the corresponding regions in the pro-GDF8 crystal structure (pdb: 5NTU), which include the prodomain α1-helix, lasso helix, fastener, and β1-strand and the growth factor β6’–β7’ strands. Prodomain monomers 1 and 2 (Pro1 and Pro2) are in cyan, and growth factor monomer 1 (GF1) is in light pink. Peptides with enhanced HDX are numbered and colored blue in Pro1 and Pro2 and magenta in GF2.

HDX reports only on backbone amide hydrogens, as deuterium incorporated at other positions such as sidechains reverts back to hydrogen during analysis (Wales & Engen, 2006). Backbone positions that are protected from HDX exchange include those that are buried and thus less accessible to solvent and those that are strongly hydrogen bonded. For comparison to GDF8 pro-complexes, we also measured HDX of a pro- TGF-β1 R249A PC-site mutant at pD 7.5, extending previous data at pD 8 (Dong, Zhao et al., 2017). Alignment of the pro-TGF-β1 and GDF8 pro-complexes enables comparison of exchange in homologous regions (Fig. 3A).

Before coming to differences among the three types of GDF8 pro-complexes, we first describe overall shared features, along with differences and similarities with pro- TGF-β1. Fig. 3A color-codes the amount of HDX of peptides after 1 min of labeling. Exchange at 5 time points between 10 s and 4 h is shown in Fig. 3B for selected peptides and for all peptides in Supplemental Figs. S1 and S2. Low deuteration in both TGF-β1 and GDF8 pro-complexes is found in the prodomain at the C-terminal portion of the α1-helix, in the β1, β3, β4, β10-strands, and in the GF in the α2’-helix and the β6’ and β7’ strands, supporting overall structural similarity. In TGF-β1, the prodomain α1-helix and β1-strand pack against the GF α2’-helix and β6’ and β7’ strands (Shi et al., 2011). Additionally, the prodomain β1-strand hydrogen bonds to the GF β7’ strand to link prodomain and GF β-sheets into a super-β-sheet. That the α1-helix, β1-strand, and β6’ and β7’ strands are among the slowest exchanging regions in TGF-β1 supports their importance in maintaining prodomain–GF interactions and hence latency. Furthermore, these are among the slowest exchanging regions in GDF8, suggesting that similar interactions between prodomain and GF elements maintain latency in TGF-β1 and GDF8 (Figs. 3A, S1 and S2).

The latency lasso in the TGF-β1 prodomain loosely wraps around the GF α2’-helix and the tip of the ‘finger’ formed by the GF β6’ and β7’ strands. The latency lasso varies among pro-TGF-β1 structures (Dong et al., 2017), and in agreement, shows fast exchange (Fig. 3A). Contrasting results were obtained with the region of GDF8 corresponding to the C-terminal portion of the latency lasso, which contains six more residues than in TGF-β1 (Figs. 3A, S1 and S2). The GDF8 latency lasso was cleaved by pepsin into two peptides. While the N-terminal GDF8 latency lasso-like peptide was highly deuterated (>60% at 1 min) like pro-TGF-β1, the C-terminal peptide was much less deuterated. Thus, we concluded that the C-terminal portion of the longer latency lasso of GDF8 must adopt a more compact, stable structure, consistent with its high content of Leu and Pro residues, and may interact with the GF and contribute to latency (Le et al., 2017). Indeed, the pro-GDF8 crystal structure shows that this insert forms a lasso α-helix that forms a hydrophobic interface with the GF finger tips (Fig. 3C) (Cotton et al., 2017).

Comparisons among the three types of GDF8 pro-complexes reveal important similarities and differences. The HDX-MS profiles of pro and latent GDF8 are essentially identical, even in the C-terminal peptide of the prodomain and N-terminal peptide of the GF that flank the PC-cleavage site (Fig. 3A). Deuterium incorporation is superimposable from 10 s to 4 h (Supplementary Fig. S1 and S2), showing that PC cleavage between the pro and GF domains has no effect on the noncovalent structure or dynamics of the pro-complex. In contrast, cleavage with TLL2 at the TLD site markedly increases deuterium exchange in regions of prodomain–GF association. The TLD-cleavage site between Arg-98 and Asp-99 in pro-GDF8 is disordered in its crystal structure (Cotton et al., 2017) and corresponds in TGF-β1 to a region that follows the α2-helix (Fig. 3A, C). Two peptides that flank the TLD-site, YDVQR^98^ and D^99^DSSDGSL, were recovered only in primed GDF8 (Fig. 3A) and confirmed TLL2-cleavage and its correlation with enhanced HDX in regions of putative prodomain–GF interactions as described in the next paragraph. However, recovery of some peptides spanning the cleavage site in primed GDF8 (residues 94-106 and 95-106, Supplementary Fig. S1) also confirmed that TLL2 cleavage was not complete.

TLD-site cleavage enhanced HDX in peptides that cluster to interacting regions of the straitjacket and GF. Based on structural alignments between GDF8 and TGF-β1, these changes in HDX correspond to the predicted α1-helix, latency lasso, α2-helix, fastener, and β1-strand in the GDF8 prodomain and the β6’ and β7’ strands in the GDF8 growth factor, as now confirmed by the pro-GDF8 crystal structure (Fig. 3A–C). Greater deuteration in these regions after TLL2 cleavage suggests structural destabilization with increased conformational dynamics and flexibility. Peptides with more exchange in primed GDF8 compared to pro-GDF8 and latent GDF8 are shown in Fig. 3C with darker colors than other segments of the prodomain and GF and are numbered identically in Figs. 3A–C as peptides 2–7 and 10. Peptides 1, 8 and 9 in Fig. 3A and B are, by comparison, peptides in the N- and C-prodomain fragment and GF that show no difference in exchange among the three types of GDF8 pro-complexes. All peptides with more HDX in primed GDF8 cluster to GF-interacting regions of the N-prodomain fragment (i.e., the C-terminal portions of the α1-helix and latency lasso), the C-prodomain fragment (i.e., the β1-strand and adjacent loops), and the GF (i.e. the finger). Thus, TLD-site cleavage increases the structural lability of these regions. Differences in HDX among peptides in a single type of pro-complex provide further structural insights. The deuterium incorporation graphs in Figs. 3B, S1 and S2 have scales in which the maximum value of the y-axis corresponds to the maximum number of backbone amide hydrogens in each peptide that can take up deuterium. Dashed lines show 50% exchange. Peptide 2 (IKIQIL) and overlapping peptide 56–66 (Fig. S1) in the C-terminal portion of the α1-helix are two of the slowest exchanging peptides in latent GDF8 (Figs. 3B, S1 and S2). These results show that the C-terminal portion of the α1-helix has a very important role in stabilizing GDF8 and further suggest that the increased α1-helix dynamics observed after TLD cleavage is likely to make an important contribution to GDF8 activation.

### EM

We used EM to define the overall shape of GDF8 pro-complexes and explore whether furin or TLL2 cleavage induced large-scale conformational change during GDF8 pro-complex maturation and activation. Pro-complexes were subjected to SEC (Fig. 4A) and immediately applied to EM grids at concentrations of 1 to 5 pM, i.e., concentrations that were substantially lower than used in SEC-MALS or HDX. Primed GDF8 eluted later than pro and latent GDF8 (Fig. 4A), as also shown in Fig. 1D, and the main peak of primed GDF8 was followed by a broad shoulder in which the GF and C-terminal prodomain fragment were prominent (Fig. 4B). The N-prodomain fragment was present in the main primed GDF8 peak and in fractions immediately after this peak including fraction 19, but was less prominent in the trailing portions of the shoulder. The C-prodomain fragment was prominent in both the peak and trailing fractions. These results suggest that some degree of dissociation of primed GDF8 occurred during gel filtration. GDF8 pro-complexes from peak fractions were electrostatically adsorbed to glow-discharged carbon grids, stained and fixed with uranyl formate, examined by electron microscopy, and ~5000 particles were subjected to alignment, classification, and averaging (Supplementary Fig. S3).

**Figure 4.**
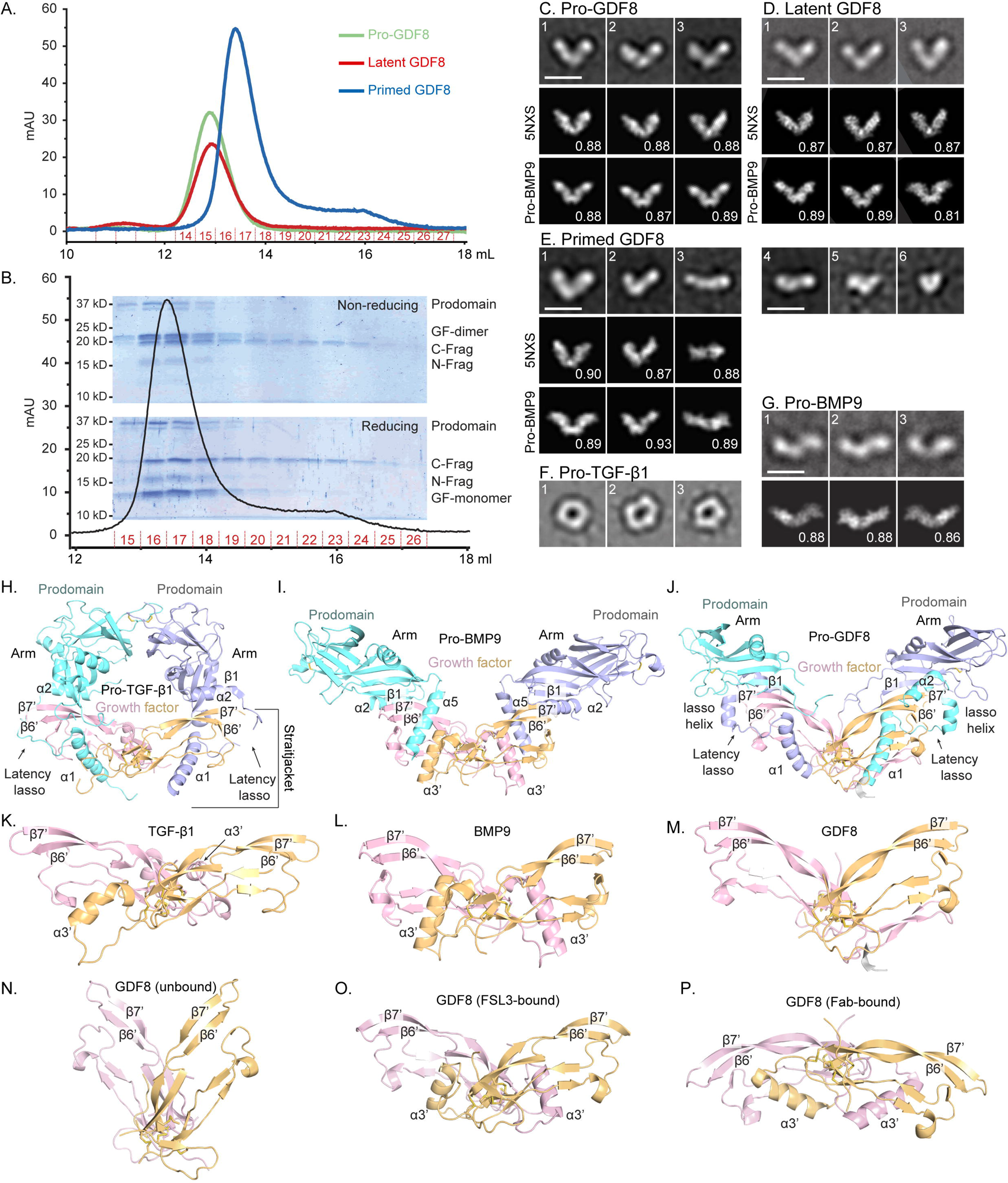
Conformation of GDF8 pro-complexes under negative stain electron microscopy (EM). **A.** Gel filtration elution profiles (S200 column) of pro-, latent, and primed GDF8 immediately prior to negative staining. B. Coomassie blue-stained SDS-PAGE gels of elution fractions 15–26 from gel filtration of primed GDF8 (A) under non-reducing (top) and reducing (bottom) conditions. **C–G**. Representative EM class averages of pro-GDF8 (C), latent GDF8 (D), primed GDF8 (E), pro-TGF-β1 (F, (Shi et al., 2011)), and pro-BMP9 (G,(Mi et al., 2015)). Scale bars, 100 Å. Class averages of GDF8 pro-complexes (C–E) were cross-correlated with 2D projections of pro-GDF8 (pdb: 5NXS) and pro-BMP9 (pdb: 4YCG) crystal structures, whereas cross-correlations of pro-BMP9 class averages with the pro-BMP9 crystal structure was previously performed in (Mi et al., 2015); the best-correlating projection and its correlation coefficient are shown below each class average. **H–J.** Crystal structures of cross-armed pro-TGF-β1 (pdb: 5FFO) (H), open-armed pro-BMP9 (pdb: 4YCG) (I), and V-armed pro-GDF8 (pdb: 5NTU) (J) superimposed on the cystine knot of the growth factor dimer. Important secondary structures involved in prodomain–GF interactions are labeled for each pro-complex structure. **K–M.** Conformation of the growth factor dimer from pro-complex crystal structures. The growth factor dimer adopts a closed conformation in pro-TGF-β1 (pdb: 3RJR) (K) (Shi et al., 2011) and pro-BMP9 (pdb: 4YCG) (L)(Mi et al., 2015), and an open conformation in pro-GDF8 (pdb: 5NTU). **N–P.** Crystal structures of the GDF8 growth factor in different conformations. The *apo* form (pdb: 5JI1)(Walker et al., 2017a) adopts an open conformation (N). In contrast, the GDF8 growth factor adopts a closed conformation when in complex with the antagonist FSL3 (O) (pdb: 3sek) (Cash, 2012) and yet another conformation when bound to Fab (P) (pdb: 5f3b) (Apgar et al., 2016).

Negative stain EM of pro- and latent GDF8 showed a V-shaped conformation, which we term V-armed (Fig. 4C, D). This conformation contrasts with the ring-like, cross-armed conformation of latent pro-TGF-β1 (Shi et al., 2011) (Fig. 4F). No differences between pro- and latent (furin-cleaved) GDF8 were detectable in EM, consistent with their essentially identical HDX (Figs. 3, S1, & S2). We have previously termed pro-BMP9 open-armed based on its EM and crystal structures (Fig. 4G) (Mi et al., 2015) and to contrast it with cross-armed pro-TGF-β1. Crystal structures show that pro-BMP9, pro-GDF8, and pro-activin A may also be considered V-armed, with a more obtuse V-angle in pro-BMP9 and pro-GDF8 (Cotton et al., 2017, Mi et al., 2015) (Fig. 4I,J) than in pro-activin A (Wang et al., 2016). Pro- and latent GDF8 class averages cross-correlate essentially equally well with the pro-GDF8 and pro-BMP9 crystal structures (bottom two rows in Fig. 4 C, D). In contrast, pro- and latent GDF8 class averages do not cross correlate as well with the pro-activin A crystal structure which has a more acute V angle (Le et al., 2017, Wang et al., 2016). SAXS also indicates that pro-GDF8 adopts an open or V-armed conformation rather than a cross-armed conformation, albeit with SAXS envelopes fitting better to the pro-activin A than the pro-BMP9 crystal structure (Walker et al., 2017b).

In EM, pro-BMP9 appears more linear than V-armed (Fig. 4G). The pro-BMP9 crystal structure projections that cross-correlate best with EM class averages of pro-GDF8 differ in orientation from those that cross-correlate best with EM class averages of pro-BMP9 (compare bottom rows of Fig. 4C, D, G). Thus, it appears that pro-BMP9 and pro-GDF8 differ in their preferred orientations on EM grids. Accordingly, the V-angle of pro-complexes in negative stain EM should be interpreted with caution, because it is dependent on orientation on the grid, which may differ among TGF-β1 family members.

In contrast to pro- and latent GDF8, EM of primed GDF8 revealed marked heterogeneity in conformation and size (Fig. 4E). Some primed GDF8 particles adopted a V-armed conformation similar to pro- and latent GDF8 (Fig. 4E, panels 1 and 2). A linear species resembled EM class averages of pro-BMP9 (compare Fig. 4E panel 3 with Fig. 4G). Class averages of progressively smaller-sized particles in Fig. 4E panels 4-6 may correspond to partially dissociated primed GDF8 containing a GF dimer and lacking one or more N and C-terminal prodomain fragments, or completely dissociated growth factor dimers or prodomain fragments. The complete class averages of the three types of pro-complexes (Supplementary Fig. S3) highlight the comparative heterogeneity of primed GDF8, which was seen with 3 independent primed GDF8 preparations. This heterogeneity was seen despite the immediate application of the main peak from SEC of primed GDF8 to EM grids. Heterogeneity seen after cleavage at the TLD-site is consistent with more rapid deuterium uptake seen with HDX-MS, the smaller molecular mass of the main peak of primed GDF in SEC-MALS, and the partial dissociation during SEC of primed GDF8 that was evident from the shoulder trailing the main peak.

## Discussion

These studies illuminate important biochemical, functional, and structural aspects of GDF8 maturation and activation. The pro-GDF8 precursor complex adopted a V-armed conformation in EM. Latent GDF8 produced by furin cleavage of pro-GDF8 adopted an essentially identical V-armed conformation. This V-armed conformation was distinct from the cross-armed conformation of latent pro-TGF-β1. Our EM results are consistent with concurrent crystal structure and SAXS studies on GDF8 pro-complexes (Cotton et al., 2017, Le et al., 2017, Walker et al., 2017b). As expected, the prodomain and GF dimer remained stably noncovalently associated in latent GDF8. Notably, HDX-MS showed indistinguishable deuteration profiles, even in peptide segments adjacent to the PC-cleavage site. Perhaps this should not be surprising, because segments that are susceptible to furin cleavage must be sufficiently flexible prior to cleavage to access the active site cleft in the large PC protease. Crystal structures of pro-TGF-β1 with and without PC cleavage and of pro-activin A with and without cleavage at an artificial, eutopic cleavage site showed only small structural changes in the immediate vicinity of the cleavage site (Wang et al., 2016, Zhao, 2017), in agreement with our EM and HDX-MS results.

Previous studies have shown that prodomain cleavage by Tolloid proteases activates GDF8 and GDF11 signaling (Ge et al., 2005, Wolfman et al., 2003); however, whether cleavage immediately released the GF from embrace by either or both of the two cleaved prodomain fragments was not examined. The issue of the fate of the prodomain fragments following Tolloid family cleavage is biologically important, because the GDF8 prodomain not only shields it from its type I and type II signaling receptors, but also from inhibitors including follistatin and follistatin-like 2, that strongly negatively regulate GDF8 signaling in vivo (Hinck et al., 2016). Thus, association with prodomain fragments could shield the GF from its inhibitors prior to or during association with its type I and type II receptors; receptor association to TGF-β1 family members can often proceed in two steps in which the GF first associates with one class of receptor and then the other (Hinck et al., 2016, Sengle, Ono et al., 2008).

Here we have described, using GDF8, a class of TGF-β1 family pro-complex we term primed, in which cleavage of the prodomain does not immediately lead to GF dissociation from prodomain fragments, but primes the GF for subsequent dissociation. Whether both the N-prodomain and C-prodomain fragments must dissociate prior to binding of the GDF8 GF to its receptors, or whether fragment dissociation is stepwise and can be coordinated with stepwise binding to receptors is an important topic for future research. We found that primed GDF8 was labile; partial N and C-terminal prodomain fragment dissociation occurred on the timescale of sample handling and gel filtration in a concentration-dependent fashion when samples were applied at concentrations of 0.13 to 1.3 µM and diluted during the gel filtration process to lower concentrations. Some dissociation occurred during gel filtration prior to EM of pM concentrations of primed GDF8,. In contrast, at the higher final concentrations used in HDX-MS, (0.7 to 1.7 µM) there was no evidence for dissociation: two or more individual peptides in the N-terminal and C-terminal prodomain fragments and the GF of primed GDF8 showed essentially identical HDX to peptides in pro and latent GDF8 (peptides 1, 7, & 8 in Fig. 3B, Figs. S1&S2).

We were unable to determine if TLL2 cleavage alone resulted in a shape change in primed GDF8 compared to pro and latent GDF8, because the small percentage (~11%) of class averages that resembled the V-shaped conformation characteristic of pro and latent GDF8 (Fig. S3) might have corresponded to the small proportion of uncleaved, latent GDF8 present in GDF8 preparations (Fig. 4B). Clearly, the affinity of the association of the GF with the prodomain, which will depend on the structural details and energetics of their association interfaces, will be more closely linked to latency than overall shape in EM. Furthermore, shape in negative stain EM is influenced by multiple factors including orientation on the substrate, as emphasized here by optimal cross-correlation of GDF8 and BMP9 pro-complex EM class averages with different projections of the BMP9 pro-complex crystal structure. In short, the apparent angle of the V in EM is influenced by orientation of the plane of the V in the pro-complex with respect to the plane of the EM grid. Another important influence on the shape in EM of TGF-β1 family pro-complexes is the orientation between the two GF monomers. In pro-activin A (Wang et al., 2016) and pro-GDF8 crystal structures (Fig. 4J, M), the GF dimer has an open conformation that contrasts with the closed GF dimer conformation in pro-complexes of TGF-β1 (Fig. 4H, K) and BMP9 (Fig. 4 I, L). If the GF in the pro-activin A complex had a closed conformation, as seen in some activin GF complex structures with inhibitors and receptors, the pro-complex would have to assume a markedly more obtuse V angle (Hinck et al., 2016, Wang et al., 2016). Notably, the apo GDF8 GF dimer structure adopts an open GF conformation (Fig. 4N) (Walker, Czepnik et al., 2017a) with a more acute V-angle than in the pro-complex (Fig. 4M), whereas crystal structures of the GDF8 GF in complex with antagonists reveal a closed conformation and a complex with Fab reveals yet another conformation (Fig. 4 O,P) (Apgar, Mader et al., 2016, Hinck et al., 2016). Our HDX results showed no changes in GDF8 GF regions that alter in conformation between the open and closed GF conformations, including the GF α3’-helix (Hinck et al., 2016), and thus provided no evidence for a change in overall GF conformation associated with TLL2 cleavage.

Prodomain cysteine residues can have important roles in the TGF-β family, and our HDX-MS studies provide information about the structural context of the four cysteine residues present in the GDF8 prodomain. At a bowtie knot at the end of the arm domain distal from the GF, pro-TGF-β contains cysteines that dimerize the prodomain and may be important for the overall cross-armed, ring-like conformation of pro-TGF-β1 (Shi et al., 2011). Mutational removal of these cysteine residues activates TGF-β1 (Brunner, Marquardt et al., 1989). Thus, while latent TGF-β1 and GDF8 may share many of the interactions between their pro and GF domains that confer latency, latent GDF8 appears to contain additional latency-conferring structural features that compensate for its lack of prodomain dimerization; the lasso α-helix may be one of these (Cotton et al., 2017). In the GDF8 prodomain, two pairs of cysteines are present in segments that align with the association region and the β2-strand of TGF-β1 (asterisks, Fig. 3A). These pairs of cysteines are separated by the TLD-cleavage site; since the N-prodomain and C-prodomain fragments are not disulfide linked to one another (Fig. 1B), cysteines in the association region do not disulfide bond to cysteines in the β2-strand. In pro-TGF-β1 the β2-strand is an edge β-strand in a β-sheet and therefore is not as constrained structurally in family evolution as middle β-strands; we earlier proposed a distinct conformation in GDF8 based on the much greater HDX of the peptide containing Cys-137 and Cys-138 in GDF8 than the corresponding peptide in TGF-β1 (Fig. 3A) and further proposed that Cys-137 and Cys-138 might disulfide bond to one another to form a vicinal disulfide bond (Le et al., 2017), which would be incompatible with β-strand conformation (Ruggles, Deker et al., 2009). Indeed, the pro-GDF8 structure demonstrates such a vicinal disulfide bond, that the β2-strand is only 2 residues long in one monomer and not formed in the other monomer, compared to 7 residues long in pro-TGF-β1, and that adjacent sequence that is part of the β2-strand in TGF-β1 is disordered in pro-GDF8 (Cotton et al., 2017).

While we cannot rule out an intramolecular disulfide bond between Cys-39 and Cys-42 in the N-prodomain fragment, the presence of these cysteines in a region that corresponds to the association region of pro-TGF-β1 leads us to suggest that they disulfide link to an as yet unidentified molecule. Cys-33 in the association region of pro-TGF-β1 becomes disulfide linked to either latent TGF-β binding proteins (LTBPs) for storage in the extracellular matrix or to glycoprotein-A repetitions predominant protein (GARP) for anchorage on the cell surface (Hinck et al., 2016). In the absence of such a partner, the association region in pro-TGF-β1 enjoys high HDX and a conformation that varies depending on the lattice environment in crystals (Dong et al., 2017). Similarly, peptides containing Cys-39 and Cys-42 in the putative association region of GDF8 show high amounts of exchange (Fig. 3A). While GDF8 pro-complexes can bind to both LTBP3 and perlecan, both interactions are noncovalent (Anderson et al., 2008, Sengle, Ono et al., 2011). Thus, we propose that yet another molecule may become disulfide linked to Cys-39 and Cys-42. If this molecule either as a monomer or dimer could disulfide link to each prodomain monomer, it would increase the avidity of the prodomain and the N-prodomain fragment for the GDF8 GF. A putative association region in activin A also contains a Cys residue, is disordered in crystal structures, and has been proposed to associate with an unidentified molecule (Wang et al., 2016).

HDX-MS comparisons of primed GDF8 to pro- and latent GDF8 provided insights into the interactions between the prodomain and GF that are weakened by TLD cleavage. HDX-MS results for regions N-terminal of the TLD-cleavage site showed cleavage-enhanced exchange in multiple overlapping peptides in each of the α1-helix, latency lasso, and α2-helix. The increased HDX in the latency lasso was especially interesting, because it occurred in a region where GDF8 has an insertion of 6 residues and markedly more hydrophobic residues compared to TGF-β1, and where HDX in GDF8 is far less than in TGF-β1 (Figs. 3, S3). The compact structure we suggested for the 6-residue insert (Le et al., 2017) corresponds to the lasso α-helix (Cotton et al., 2017). TLD cleavage also enhanced exchange of peptides in regions C-terminal to the cleavage site that correspond to the fastener and β1-strand. The Tyr-Tyr dipeptide fastener sequence in TGF-β1 is conserved as a Tyr-His sequence in GDF8 (Fig. 3A), and both interact with the C-terminal portion of the α1-helix to secure the straitjacket (Cotton et al., 2017, Shi et al., 2011). In TGF-β1, the fastener resists TGF-β1 activation by force (Dong et al., 2017).

Besides these regions of enhanced exchange, which were proximal in sequence to the cleavage site and correspond to all major elements of the straitjacket plus the β1-strand of the arm domain, the only other peptides in primed GDF8 that showed a marked increase in exchange were two overlapping peptides that are ~300 residues C-terminal to the cleavage site and cover the GF β6’ and β7’-strands. These results provide strong evidence that straitjacket cleavage results in enhanced HDX, with increased exposure of the peptide backbone to solvent or increased dynamics of the peptide backbone, and that straitjacket backbone perturbation is transmitted to the backbones of the GF β6’ and β7’-strands.

Alterations in HDX-MS in primed GDF8 suggest that straitjacket elements including the α1-helix, latency lasso, and α2-helix and arm domain β1-strand interact with one another and with the GF β6’ and β7’-strands in GDF8 in a manner very similar to that revealed in the structure of pro-TGF-β1 (Shi et al., 2011). The pro-GDF8 crystal structure (Cotton et al., 2017) further supports similarities in straitjacket–-GF interactions between GDF8 and TGF-β1 pro-complexes. In these structures, the arm domain β1-strand hydrogen bonds to the GF β7’-strand to link arm domain and GF β-sheets into a super β-sheet. The straitjacket α2-helix covers the super β-sheet junction. The latency loop wraps around the GF β6’ and β7’-strands that form two GF fingers. The straitjacket fastener links to the α1-helix to encircle the GF fingers on the end opposite from the latency lasso (Fig. 3C).

Our results provide a compelling model for the mechanism by which TLL2 cleavage primes GDF8, i.e. releases GDF8 from latency. Many of the regions that are most strongly protected from exchange with solvent in the structure of latent GDF8 become available for exchange after cleavage at the TLD site. Regions that have low HDX are more structurally stable; thus, the greater rate of exchange of the GF straitjacket, arm β1-strand, and the interacting GF β6’ and β7’-strands in primed GDF8 can be directly related to structural destabilization with greater exposure to solvent, lower affinity between the prodomain and the GF, and dissociation of the prodomain fragments from the GF.

Our results are also in excellent agreement with mapping of inhibitory prodomain fragments (Jiang, Liang et al., 2004, Ohsawa, Takayama et al., 2015, Takayama, Noguchi et al., 2015). The size of the minimum fragment found to be required for inhibition varied among the studies and may have correlated inversely with the concentration of the inhibitors that were achieved. Prodomain fragments used as inhibitors were derived either by addition of purified bacterial fusion proteins, by co-transfection of mammalian cells with Fc fusions, or by addition of purified peptides. Using these three methods for obtaining inhibitory fragments, respectively, the minimal inhibitory fragment was found to include the entire straitjacket plus the arm domain β1-strand (Jiang et al., 2004), half of the association region, the α1-helix, and the latency lasso (Ohsawa et al., 2015), or a 23-residue α1-helix peptide (Takayama et al., 2015). The concentration required for half-maximal inhibition by the peptide, ~10 μM, was far higher than for the intact prodomain, 1 nM (Thies et al., 2001). Nonetheless, our findings that the straitjacket α1-helix has the slowest HDX of the entire GDF8 straitjacket, and that the entire straitjacket region plus the arm domain β1-strand show increased exchange with solvent in primed GDF8, are in excellent agreement with both the most minimal fragment and the largest fragment found in these studies, respectively.

In summary, our study has demonstrated that Tolloid cleavage does not immediately result in release of the GDF8 GF, but primes it for release from prodomain fragments. The HDX dynamics of GDF8 provide insights into pro-complex structure and identify a cluster of interacting structural elements that are buried in the prodomain complex with the GF. Tolloid cleavage weakens these interactions and primes the GF for subsequent release from inhibitory embrace by the prodomain.

## Materials and Methods

### Expression and purification of proteins

Stable cell lines overexpressing N-terminally 6x His-tagged pro-GDF8 (accession: O14793), a C-terminally FLAG and 6X His-tagged human furin construct (residues 1-595) (accession: P09958), and a C-terminally FLAG and 6X His-tagged full-length human TLL2 construct (accession: Q9Y6L7) were established by stable integration of plasmids in Flp-In T-REX 293 cells (Life Technologies, Carlsbad, CA). Cell lines were adapted to suspension growth in F17 media (Life Technologies, Carlsbad, CA) and proteins were expressed according to manufacturer’s instructions.

Different GDF8 pro-complexes were prepared as follows. Pro-GDF8 was purified from the supernatants of pro-GDF8 transfectants cultured in the presence of 30 μM decanoyl-Arg-Val-Lys-Arg-choloromethylketone (R&D Systems). Latent GDF8 was produced via in vitro cleavage of purified pro-GDF8 by human furin protease, which had been purified by Ni-NTA chromatography. Primed GDF8 was produced by in vitro cleavage of purified pro-GDF8 utilizing conditioned media from stable TLL2 expressing cells and purified furin protease. All in vitro cleavage reactions occurred at 30°C for 24 hours, in protease cleavage buffer: 100 mM HEPES pH 7.5, 0.01% Brij-35, 1 mM CaCl2, and 1 μM ZnCl_2_.

After five days of expression, culture supernatant was collected and cleared by centrifugation for 10 minutes at 450 x gravity at 4°C. Supernatant was then filtered by passing it through a 0.22 µm pore filter. Filtered supernatant was combined with Tris, NaCl and NiCl_2_ (final concentration of 50 mM Tris pH 8.0, 350 mM NaCl, and 0.5 mM NiCl_2_), purified by Ni-NTA chromatography (Qiagen) in 20 mM Tris, pH 8.0, 500 mM NaCl and 20 mM imidazole, and eluted with 20 mM Tris, pH 8.0, 500 mM NaCl, and 300 mM imidazole. The protein was further purified by Superdex 200 size exclusion chromatography (SEC) equilibrated with 20 mM HEPES pH 7.5, 150 mM NaCl. An additional FLAG resin purification step (GenScript, proceeded according to manufacturer’s directions) was applied as needed for removal of the FLAG-tagged proteases from the latent and primed preparations of GDF8. Peak fractions were pooled and concentrated to 1–2 mg/mL and aliquots flash-frozen and stored at -80°C.

For insect cell expression, full-length pro-GDF8 N-terminally tagged with His-SBP was cloned into the S2-2 vector (ExpreS2ion Biotechnologies) and stably integrated into *Drosophila* S2 cells. Cells were adapted to growth in serum-free Excell 420 media. After 4 days, culture supernatant was collected, filtered, buffer exchanged to 20mM Tris-HCl pH 7.5, 500 mM NaCl, and purified by Ni-NTA chromatrography as described above. After a Precision3C-cleavage step to remove the His-SBP tag, pro-GDF8 was dialyzed against 20mM Tris-HCl pH 7.5, 150 mM NaCl and subjected to another round of Ni-NTA chromatography followed by Superdex 200 size exclusion chromatography (SEC) equilibrated with 20mM Tris-HCl pH 7.5, 150 mM NaCl.

### Size exclusion and light scattering

Analysis of human pro-GDF8, latent GDF8, and primed GDF8 by size exclusion chromatography coupled to multi-angle light scattering was performed by the Keck Biophysics Facility at Yale University. Samples were analyzed at concentrations of 1.3 μM, 0.26 μM, and 0.13 μM (primed only). The light scattering data were collected using a Superdex 200, 10/300, HR Size Exclusion Chromatography (SEC) column (GE Healthcare, Piscataway, NJ), connected to High Performance Liquid Chromatography System (HPLC), Agilent 1200, (Agilent Technologies, Wilmington, DE) equipped with an autosampler. The elution from SEC was monitored by a photodiode array (PDA) UV/VIS detector (Agilent Technologies, Wilmington, DE), differential refractometer (OPTI-Lab rEx Wyatt Corp., Santa Barbara, CA), static and dynamic, multiangle laser light scattering (LS) detector (HELEOS II with QELS capability, Wyatt Corp., Santa Barbara, CA). The SEC-UV/LS/RI system was equilibrated in buffer (20 mM HEPES pH 7.5, 150 mM NaCl) at the flow rate of 0.5 ml/min or 1.0 ml/min. Two software packages were used for data collection and analysis: the Chemstation software (Agilent Technologies, Wilmington, DE) controlled the HPLC operation and data collection from the multi-wavelength UV/VIS detector, while the ASTRA software (Wyatt Corp., Santa Barbara, CA) collected data from the refractive index detector, the light scattering detectors, and recorded the UV trace at 280 nm sent from the PDA detector. The weight average molecular masses, Mw, were determined across the entire elution profile in the intervals of 1 sec from static LS measurement using ASTRA software as previously described (Folta-Stogniew & Williams). For each set of data, the level of glycosylation was established from the “Three Detector” approach.

### Reporter cell assays

Samples of pro-GDF8, latent GDF8, primed GDF8, and mature GDF8 growth factor (R&D systems) were incubated at different concentrations with 293T cells containing a stably integrated pGL4 plasmid (Promega, Madison, WI) with a promoter comprising 12 repeats of the SMAD-responsive CAGA sequence (AGCAGACA) (Thies et al., 2001). Cells were incubated at 37°C for 6 hours before detection of luciferase expression using BRIGHT-GLO^TM^ reagent (Promega, Madison, WI) according to manufacturer’s instructions. EC50 values were calculated from four technical replicates in Prism 7.01 using a variable slope four parameter non-linear curve fit.

### Hydrogen-deuterium exchange mass spectrometry

HDX-MS experiments were performed using methods reported previously (Iacob, 2013). The three GDF8 pro-complex forms were analyzed as follows. 3 µL of pro-GDF8 (25.4 µM), latent GDF8 (11 µM), and primed GDF8 (20 µM) were individually diluted 15-fold into 20 mM Tris, 150 mM NaCl, 99% D_2_O (pD 7.5) at room temperature for deuterium labeling. At time points ranging from 10 sec to 240 min, an aliquot was taken and deuterium exchange was quenched by adjusting the pH to 2.5 with an equal volume of cold 150 mM potassium phosphate, 0.5 M tris (2-carboxyethyl) phosphine hydrochloride (TCEP-HCl), H_2_O. Each sample was analyzed as previously described (Iacob, 2013, Wales, 2008). Briefly, the samples were digested online using a Poroszyme immobilized pepsin cartridge (2.1 mm x 30 mm, Applied Biosystems) at 15 °C for 30 s, then injected into a custom Waters nanoACQUITY UPLC HDX Manager^TM^ and analyzed on a XEVO G2 mass spectrometer (Waters Corp., USA). The average amount of back-exchange using this experimental setup was 20–30%, based on analysis of highly deuterated peptide standards. All comparison experiments were done under identical experimental conditions such that deuterium levels were not corrected for back-exchange and are therefore reported as relative (Wales & Engen, 2006). All experiments were performed in triplicate. The error of measuring the mass of each peptide averaged ± 0.12 Da in this experimental setup. The HDX-MS data were processed using DynamX 3.0 (Waters Corp., USA).

### Negative-Stain EM

Pro-GDF8, latent GDF8, and primed GDF8 were purified by SEC using Superdex 200 HR pre-equilibrated with 20 mM HEPES, pH 7.5, and 150 mM NaCl. The peak fraction was loaded onto glow-discharged carbon-coated grids, buffer was wicked off, and grids were immediately stained with 0.75% (wt/vol) uranyl formate and imaged with an FEI Tecnai T12 microscope and Gatan 4K×4K CCD camera at 52,000× magnification (2.13 Å pixel size at specimen level) with a defocus of −1.5 μm. Well-separated particles were interactively picked using EMAN2 (Rees, Langley et al., 2013). Class averages were calculated by multireference alignment followed by K-means clustering using SPIDER (Chen, Xie et al., 2010, Frank, Radermacher et al., 1996, Mi, Lu et al., 2011). Cross-correlations were with 2D projections generated at 4° intervals from 20-Å-filtered pro-GDF8 (pdb: 5NTU and 5NXS) and pro-BMP9 (pdb: 4YCG and 4YCI) crystal structures. We show only cross-correlations against 5NTU for pro-GDF8 and 4YCI for pro-BMP9 in Figure 4 because 5NXS and 4YCG yielded essentially identical results, respectively.

## Acknowledgements

The authors would like to thank Melissa Chambers and Zongli Li for help with EM data collection, Prof. Thomas Wales for insightful discussions on data processing, and Margaret Nielsen for her assistance in designing figures. We acknowledge a research collaboration with the Waters Corporation (JRE). This work was supported by NIH Grant R01CA210920. Y.T. was supported by a Komen postdoctoral research fellowship (Komen PDF15334161).

